# The distribution of fitness effects of plasmid pOXA-48 in clinical enterobacteria

**DOI:** 10.1101/2023.07.11.548518

**Authors:** Ariadna Fernandez-Calvet, Laura Toribio-Celestino, Aida Alonso-del Valle, Jorge Sastre-Dominguez, Paula Valdes-Chiara, Alvaro San Millan, Javier DelaFuente

## Abstract

Antimicrobial resistance (AMR) in bacteria is a major public health problem. The main route for AMR acquisition in clinically important bacteria is the horizontal transfer of plasmids carrying resistance genes. AMR plasmids allow bacteria to survive antibiotics, but they also entail physiological alterations in the host cell. Multiple studies over the last years indicate that these alterations can translate into a fitness cost when antibiotics are absent. However, due to technical limitations, most of these studies are based on analysing new associations between plasmids and bacteria generated *in vitro*, and we know very little about the effects of plasmids in their native bacterial hosts. In this study, we used a CRISPR-Cas9-tool to selectively cure plasmids from clinical enterobacteria to overcome this limitation. Using this approach, we were able to study the fitness effects of the carbapenem resistance plasmid pOXA-48 in 35 pOXA-48-carrying isolates recovered from hospitalised patients. Our results revealed that pOXA-48 produces variable effects across the collection of wild type enterobacterial strains naturally carrying the plasmid, ranging from fitness costs to fitness benefits. Importantly, the plasmid was only associated with a significant fitness reduction in 4 out of 35 clones, and produced no significant changes in fitness in the great majority of isolates. Our results suggest that plasmids produce neutral fitness effects in most native bacterial hosts, helping to explain the great prevalence of plasmids in natural microbial communities.

## Introduction

Plasmids are extrachromosomal genetic elements able to spread horizontally between bacteria. Plasmids contain genes responsible for their own replication, partition and transfer, as well as accessory genes that can provide a selective advantage to the host bacterium. Crucially, plasmids fuel bacterial evolution by disseminating accessory genes across populations^1, 2^. Antimicrobial resistance (AMR) is one of the most illustrative examples of the ability of plasmids to promote bacterial adaptation^3^. AMR is a major public health issue^4^, and plasmids are the most important vehicle for the dissemination of AMR genes across bacterial pathogens^5^. AMR is particularly concerning in clinical settings, where nosocomial pathogens carrying AMR plasmids represent a great threat to hospitalised patients^4, 6^.

Despite the selective advantage that plasmids may provide, they also produce physiological alterations in the bacterial host that can translate into a fitness cost^7–13^. These plasmid-associated costs raise the paradox of how plasmids are maintained in bacterial populations^14^. Specifically, in the absence of selective pressure for plasmid-encoded traits, purifying selection should remove plasmid-carrying bacteria, while in the presence of selection for plasmid-encoded genes, these loci would eventually be captured by the chromosome, making the plasmid redundant. Different solutions have been proposed for this plasmid paradox (reviewed in ^15^). A simple solution is that plasmid fitness effects differ across different bacterial hosts, and some strains experience no costs at all. We recently showed that this is indeed the case for pOXA-48^16^, a conjugative plasmid conferring carbapenem resistance, which is widely distributed across nosocomial enterobacteria worldwide^17^. In a previous study, we introduced pOXA-48 into a collection of “ecologically compatible” clinical enterobacteria^16^. We defined these isolates as ecologically compatible with the plasmid because they were recovered from hospitalised patients that although were not colonised by pOXA-48-carrying enterobacteria, coexisted in hospital wards with patients colonised with pOXA-48- carrying enterobacteria. The plasmid produced a 3% average fitness reduction in this collection; however, we observed a wide distribution of fitness effects across strains, including fitness advantages in some of them.

One general limitation in the study of plasmid fitness effects is that most of the works to date study the effects of plasmids that are introduced *de novo* in naive bacterial strains. Therefore, there is little information about the fitness effects of plasmids in the natural bacterial hosts that actually carry them^18, 19^. The reason underlying this limitation is the lack of proper methods for selectively eliminating (curing) plasmids from wild-type bacteria without producing further genetic alterations. In this study, we used a CRISPR- Cas9-based method that we recently developed^20^ to specifically cure a collection of pOXA-48-carrying enterobacteria that were recovered from hospitalised patients. With this method, we produced 35 isogenic pairs of wild-type and pOXA-48-free strains that allowed us to precisely estimate pOXA-48 fitness effects.

## Results

### Curing pOXA-48 from wild-type clinical enterobacteria

In this study, we aimed to determine the distribution of pOXA-48 fitness effects in wild-type strains naturally carrying this plasmid (the outline of this study is presented in Figure 1). We recently performed a genomic characterization of a large collection of 225 pOXA- 48-carrying enterobacteria recovered from patients in a large hospital in Madrid, Spain^21^ (Figure 2A; Table S1). In this work, we randomly selected a subset of 60 strains from that collection (Figure 2B; Table S1). We were able to transform our curing vector (pLC10) in 44 of them, and we successfully cured 35 clones belonging to 5 different species (Figure 2C,D; Table S1; see Methods and Figure S1 for details about the curing protocol).

**Figure 1.**
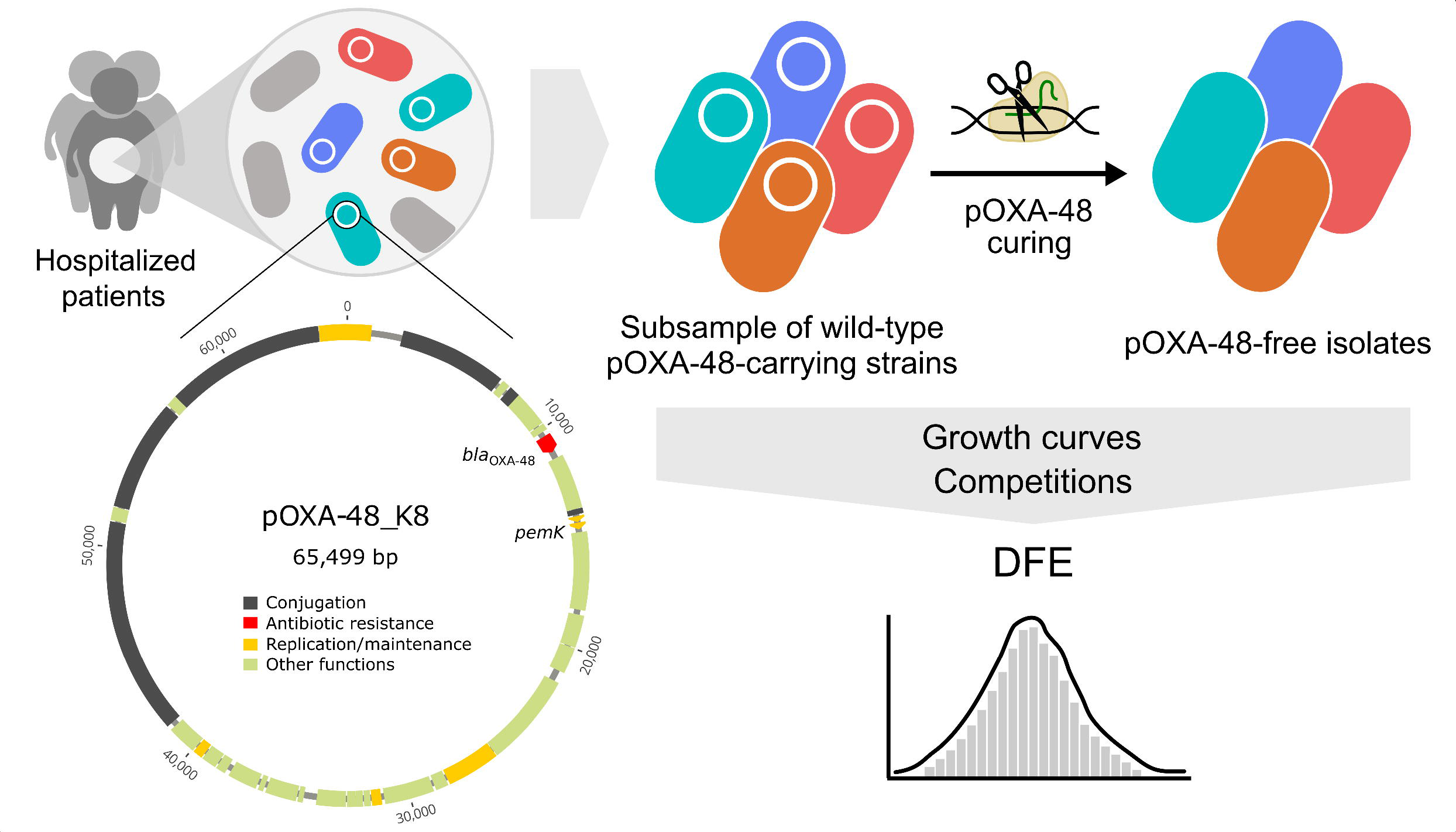
Experimental design. A representative subsample of wild-type pOXA-48- carrying enterobacteria (n = 60) was selected from a collection isolated from the gut microbiota of hospitalised patients (n = 225). Plasmid pOXA-48 was eliminated (cured) from the wild-type strains using a CRISPR-Cas9-based method (see Methods). Growth curves and competition experiments were performed for all the pOXA-48-carrying and pOXA-48-free clones. We used the relative fitness data obtained from the competition assays to determine the distribution of fitness effects (DFE) of pOXA-48 in its natural hosts.

**Figure 2.**
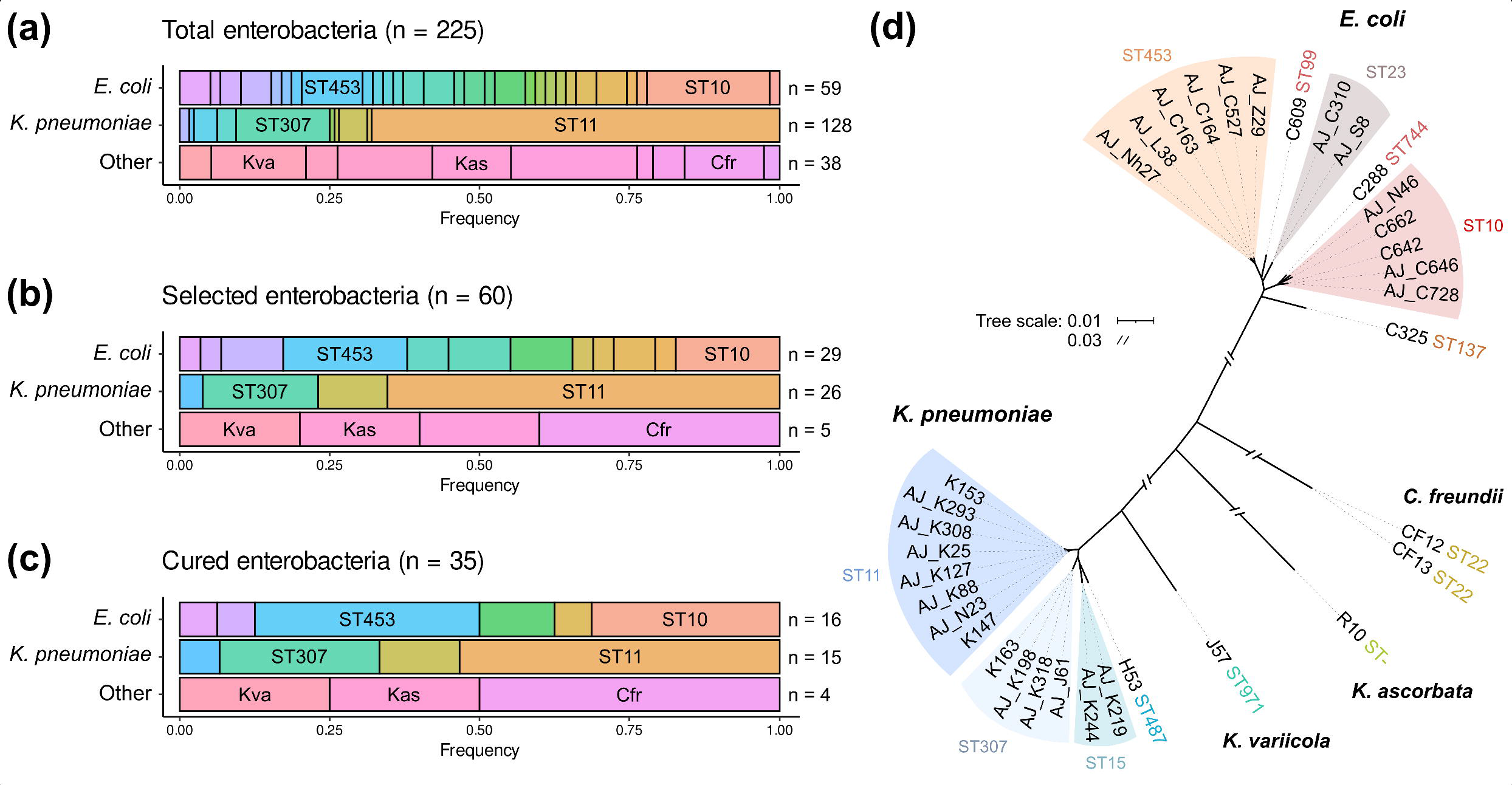
Bacterial collection used in this study. Summary of pOXA-48-carrying strains present in the entire collection and the subset used for this study. (a) Distribution of Multilocus Sequence Types (ST) for *E. coli*, *K. pneumoniae*, and other species in the collection of wild-type pOXA-48-carrying enterobacteria. (b) Representative subsample of the collection selected for this study. (c) Strains successfully cured from pOXA-48. Colours indicate different ST (upper and central rows) or different species (lower row). The most representative ST in *E. coli* and *K. pneumoniae* are indicated. Kva: *Klebsiella variicola*; Cfr: *Citrobacter freundii*; Kas: *Kluyvera ascorbata*. (d) Unrooted phylogeny constructed with the genomic sequences of the 35 strains cured from pOXA-48. Branch length represents mash distances between whole-genome assemblies. Scale breaks were introduced for long branches. The ST are also indicated.

### Genomic validation of the plasmid curing system

To validate the use of our CRISPR-Cas9 system to selectively cure pOXA-48, we sequenced and analysed in detail the genomes of nine cured strains. These included two *E. coli* (C288 and C325), two *Citrobacter freundii* (CF12 and CF13), one *K. variicola* (J57), and four *K. pneumoniae* (H53, K147, K153 and K163). The genomes of their respective wild-type strains were sequenced using long-read technology. These sequences were combined with the short-read sequences obtained previously^21^ to generate closed and complete genome assemblies to use as reference. By comparing the genomes of cured and wild-type clones, we were able to confirm that only plasmid pOXA-48 was eliminated during plasmid curing (Table S2).

To test if our system was able to generate near-isogenic plasmid-carrying and plasmid- free clones, we analysed genomic variants in the cured strains compared to the wild- type strains (Table S3). Only two strains, CF13c1 and K163c1, did not accumulate any mutations at all. The remaining seven strains presented between one and three single nucleotide polymorphisms (SNPs). Additionally, two of the strains, J57c1 and K153c2, showed possible genomic rearrangements. For a detailed description of the mutations see Supplemental Results.

Next, we investigated if the observed SNPs could be caused by off-target activity of the CRISPR-Cas9 system. We compared the sequences of the single-guide RNAs (sgRNAs) used to cure pOXA-48 with the mutated regions and found that sgRNAs aligned poorly (see Supplemental Results for more details). Thus, it is unlikely that the mutations are the result of off-target effects of the curing system, and they probably accumulated randomly due to single cell bottlenecks during the curing process. In summary, despite accumulating a few mutations, we show that wild-type and cured strains are isogenic or near-isogenic. We therefore conclude that the CRISPR-Cas9 system can be reliably used to cure pOXA-48 from clinical enterobacterial isolates.

### Determination of pOXA-48 fitness effects

We performed competition assays to determine the fitness effects of plasmid pOXA-48 in the different strains of the collection (Figure 3). We used flow cytometry to increase the throughput of these assays. Specifically, we competed each pOXA-48-carrying and pOXA-48-free clones against a fluorescently labelled *E. coli* J53 strain (n = 70). We previously showed that this strain can be used as a common competitor against wild- type enterobacteria, producing comparable results to those from competitions between isogenic clones^16^. Moreover, as a control for the competition assays, we also performed individual growth curves of every clone in the collection (n = 70). As expected, we observed a positive correlation between the relative fitness of the different clones measured in the competitions assays against *E. coli* J53 and the area under the growth curves (AUC). AUC compiles different growth curve parameters^16, 22^ (lag time, OD_max_ and V_max_) and can be used as a proxy of bacterial fitness (Spearman’s rank correlation, *rho* = 0.626, *S* = 20446, *P* = 1.84 x 10^-8^; Figure S2). Finally, we designed the competition assays to avoid the conjugation of plasmid pOXA-48 between the competitors, and we experimentally tested that neither conjugation, nor plasmid loss, affected the assay results (see Methods and Figure S3).

**Figure 3.**
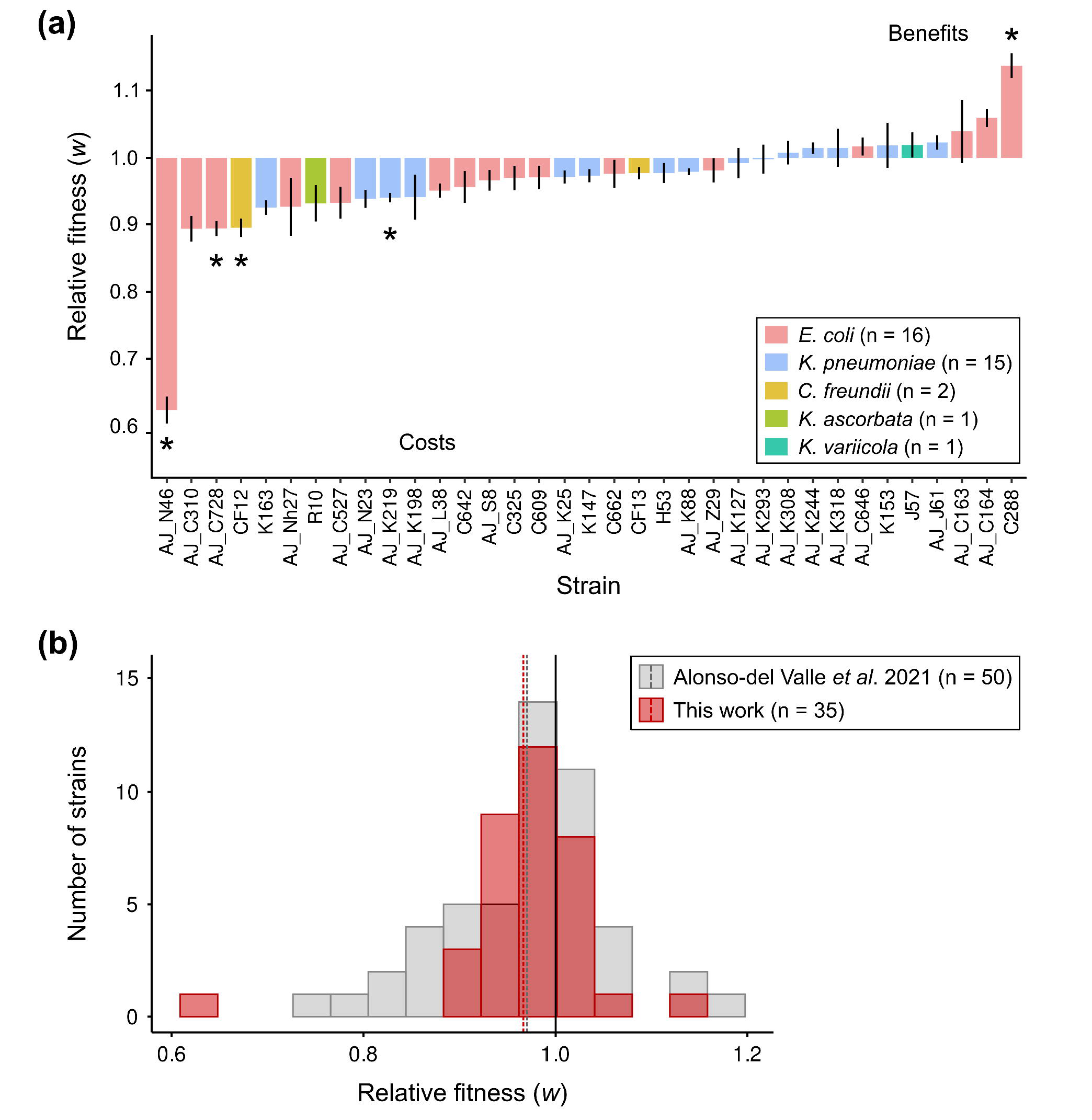
Fitness effects of plasmid pOXA-48 in wild-type clinical enterobacteria. (a) Relative fitness (*w*) of pOXA-48-carrying clones compared to their corresponding pOXA- 48-free isogenic versions (see Methods). Values above 1 indicate that plasmid pOXA-48 is associated with a fitness advantage, while values below 1 indicate a fitness reduction associated with pOXA-48 carriage. Bars represent normalised relative fitness (average of six independent biological replicates of the competition pOXA-48-carrying *vs*. *E. coli* J53 divided by the average of six independent biological replicates of the competition pOXA-48-free *vs*. *E. coli* J53). Error bars represent the propagated standard error. Asterisks indicate significant costs or benefits associated with pOXA-48 carriage (Bonferroni corrected paired *t* tests, *P* < 0.05). Strain AJ_N46 is an outlier (Grubbs test, *P* = 1.07 x 10^-6^). (b) Comparison of pOXA-48 DFE in the collection of naive, ecologically compatible pOXA-48 carriers (Alonso-del Valle *et al*. 2021^7^) and in the collection of pOXA-48 natural carriers analysed in this work. Bars indicate the number of strains in each relative fitness bin. Dashed lines indicate the mean relative fitness of each collection. Both distributions are normal when not including the outlier AJ_N46 (Shapiro-Wilk normality tests: Alonso-del Valle *et al*. 2021, *P* = 0.14; this work, *P* = 0.095). Cumulative distribution functions for both collections are represented in Figure S4.

We used the data from the competition assays to calculate the relative fitness of each wild-type pOXA-48-carrying clone compared to its pOXA-48-free isogenic version (Figure 3A). Results showed that pOXA-48 produced a range of fitness effects, spanning from costs to benefits. Specifically, plasmid carriage was associated with a significant fitness cost in four strains and produced a significant fitness advantage in one strain (Bonferroni corrected paired *t* tests, *P* < 0.05). In the remaining 30 strains, pOXA-48 produced no significant change in relative fitness. Finally, and in line with our previous results^16^, we observed no differences in the distribution of pOXA-48-induced fitness effects between *E. coli* and *K. pneumoniae* strains (Wilcoxon rank sum test, *W* = 94, *P* = 0.32; Figure S4).

### Distribution of pOXA-48 fitness effects

We previously determined the DFE of pOXA-48 in naive but ecologically compatible enterobacteria from the gut microbiota of hospitalised patients. The plasmid produced variable fitness effects with a small fitness reduction on average (mean *w* = 0.971, var = 0.0072; Figure 3B). Here, we reported the DFE in enterobacterial strains naturally carrying pOXA-48 recovered from patients in the same hospital (mean *w* = 0.967, var = 0.006; Figure 3B). Our data revealed that pOXA-48 produced similar fitness effects in enterobacterial strains that were recovered with or without the plasmid from the same cohort of hospitalised patients (Figure S5, two sample *t* test, *t* = 0.23, df = 77.67, *P* = 0.83; exact two-sample Kolmogorov-Smirnov test, *D* = 0.17, *P* = 0.56). Interestingly, our studies revealed that pOXA-48-carriage is not associated with fitness costs in the majority of wild-type enterobacterial strains from the gut microbiota of hospitalised patients, and this claim holds true both for clones naturally carrying pOXA-48 (31/35) and for pOXA-48-free clones (36/50).

## Discussion

The DFE of mutations is a central concept in genetics and evolutionary biology, with implications ranging from population adaptation rates to complex human diseases^23^. The fitness effects of new spontaneous mutations in bacteria follow a heavy-tailed distribution dominated by quasi-neutral mutations with infrequent strongly deleterious mutations^24, 25^. Horizontally acquired genes can also impose a fitness cost in bacterial hosts^26, 27^. However, horizontal gene transfer in bacteria is frequently mediated by mobile genetic elements, such as plasmids, that carry multiple genes. Over the last years, many studies have quantified plasmid-associated fitness effects^7, 10, 28–33^. Nevertheless, most of our understanding of plasmid fitness costs comes from analysing the effects of foreign plasmids artificially introduced into new bacterial hosts, and we still know very little about the effects of plasmids in their native bacterial hosts. The difficulty of selectively removing plasmids from wild-type bacteria is responsible for this important limitation. However, recent advances in genome editing have allowed us to overcome this problem. Utilising a previously developed CRISPR-Cas9-based curing system^20^, we are now able to remove the carbapenem resistance plasmid pOXA-48 from wild-type, multidrug resistant, clinical enterobacterial strains. This system presents both high efficiency (we were able to successfully cure 35 out of 44 clones transformed with the curing vector) and high specificity (no off-target effects were observed in the genomic analysis of the cured strains). Crucially, this curing system could be easily re-coded to remove virtually any plasmid from enterobacteria.

In this study, we determined the DFE of pOXA-48 in pOXA-48-carrying enterobacteria recovered from a cohort of hospitalised patients. Our findings reveal that the distribution of plasmid fitness effects in native hosts did not differ from that previously observed in ecologically compatible pOXA-48-free hosts from the same collection. The DFE was dominated by quasi-neutral mutations, with a slight shift towards fitness costs (Figure 3B). Interestingly, this DFE produced by a mobile genetic element is similar to that of spontaneous mutations. It is important to mention that in our studies we divided the clinical isolates between pOXA-48-carrying or pOXA-48-free. However, it is impossible to rule out the possibility of the previous presence of this plasmid in the pOXA-48-free isolates. The similarity of pOXA-48 effects in both collections could therefore be the result of plasmid-bacterium preadaptations driven by the high mobility of pOXA-48-like plasmids in gut microbiota communities^21^. Alternatively, the DFE observed in the collection of pOXA-48-free wild-type strains could be explained by the high permissiveness to plasmid acquisition in this collection^29^. Regardless of the underlying mechanism, our results highlight the relevance of the “ecological compatibility” between plasmids and their bacterial hosts. Supporting this idea, the costs observed in these two studies are lower than those observed in associations between plasmids and bacteria from different ecological origins (data form meta-analysis of 50 plasmid–bacterium pairs from 16 independent studies^28^, mean *w* = 0.91, var = 0.029).

The results from this and other recent studies help to explain the high prevalence of pOXA-48-like plasmids in clinical environments^17^. First, pOXA-48 produces no or moderate fitness costs in most of the enterobacterial clones in the gut microbiota of hospitalised patients^16^. Second, pOXA-48 spreads through conjugation at high frequencies in these communities, allowing the plasmid to explore new bacterial hosts^21^. And third, pOXA-48-bacteria associations experience rapid within-patient evolution, promoting their adaptation to different antibiotic regimes^20^. We argue that the different eco-evolutionary dynamics that help resolve the “plasmid paradox” for pOXA-48 will probably apply to many other plasmids across the wide diversity of natural microbiota.

## Methods

### Strains, media and culture conditions

Bacterial clones carrying pOXA-48-like plasmids included in this study were initially isolated and characterised as part of a surveillance screening for detecting extended spectrum β-lactamases/carbapenemases in hospitalised patients in the Hospital Universitario Ramon y Cajal (Madrid, Spain) (R-GNOSIS-FP7-HEALTH-F3-2011- 282512; http://r-gnosis.eu/). All clones included in this study were recovered as part of a project approved by the Hospital Universitario Ramon y Cajal Ethics Committee (ref. no. 251/13). Further details of R-GNOSIS isolation protocol and sample characterization can be found^20, 21, 34^. To study the distribution of plasmid-associated fitness effects in wild-type strains naturally carrying pOXA-48-like plasmids, a representative subset of 60 clones which included the most common sequence types in the collection (determined by Multilocus Sequence Types) was randomly generated (Table S1). Note that the most predominant STs in the collection were maintained in the final selection (n=35), but not the final proportions of each ST from the initial collection (n=225) as shown in Figure 2 (A, B, C) and Table S1. For antimicrobial susceptibility testing, Mueller Hinton II broth (MH, Oxoid) was used. Lennox lysogeny broth (LB) was used for plasmid curing, growth curves and competition assays. Chromogenic agar (HiCrome UTI Agar, HIMEDIA) was used for plasmid curing. When indicated, solid media was obtained by supplementing each medium with 15 g/L agar (CONDA). Amoxicillin-clavulanic acid (Sandoz), ertapenem, chloramphenicol, apramycin, kanamycin (Merck) were used in this study. pLC10 purification was performed with Plasmid EasyPure (Macherey-Nagel).

### pOXA-48 curing

The synthetic pLC10 plasmid carrying the CRISPR-Cas9 machinery, was used to cure the native pOXA-48 plasmid from the wild-type enterobacteria. Two different pLC10 versions were used depending on the resistance profile (Table S1) of each bacterium: pLC10-Kan (kanamycin-resistant) and pLC10-Apra (apramycin-resistant). A full description of the plasmid can be found in Figure S1 and in reference^20^. All primers used in this study are listed in Table S4. Importantly, pLC10 codes for a thermosensitive replication initiation protein (pSC101-based) which is not functional at 37°C. Cas9 protein expression is under the control of a P*_tet_* promoter which is inducible with anhydrotetracycline (ATC). Two different single-guide RNA (sgRNA) were used targeting two different pOXA-48 genes (*pemK* or *bla*_OXA-48_) and differences in curing efficiency depending on the sgRNA were not detected. The sgRNA expression is under the control of a P*_lac_* promoter, inducible with isopropyl β-d-1-thiogalactopyranoside (IPTG). Each sgRNA was introduced by Golden Gate assembly (New England Biolabs). Primers used for cloning each sgRNA were (i) for the *pemK* region, Fw 5’- CACAGTTGTGCCCGTGACCAGCGG-3’ and Rev 5’- AAACCCGCTGGTCACGGGCACAAC-3’ and (ii) for *bla*_OXA-48_, Fw 5’- CACATGGCTTGTTTGACAATACGC-3’ and Rev 5’-AAACGCGTATTGTCAAACAAGCCA-3’. Later, pOXA-48-carrying strains were made competent: we cultured each clone overnight at 37°C with continuous agitation (250 r.p.m.; MaxQ 8000, Thermo Fisher Scientific). Then, overnight cultures were diluted 1:100 in LB and after 2.5 hours of culturing in the same conditions as the day before, cells were harvested by centrifugation (3,000 g). Then, bacterial cells were washed with distilled ice-cold water (Invitrogen) several times. Later, pLC10 was introduced into bacterial cells by electroporation using 0.1 cm cuvettes / 1.8 kV pulse following manufacturer recommendations (MicroPulser Electroporator; Bio-Rad Laboratories). Transformants were selected on LB agar plates supplemented with kanamycin 250– 512 µg/mL or apramycin 30-50 µg/mL, depending on the plasmid version. pLC10 presence was determined by PCR (Fw 5’-CTCGGTAGTGGGATACGACGA-3’ and Rev 5’-CACTGAAAGCACAGCGGCTG-3’; amplicon size 859 bp). Then, CRISPR-Cas9 machinery was induced by resuspending transformants biomass in 500 µL of LB supplemented with kanamycin or apramycin and 0.3 µg/mL of ATC to induce Cas9 expression, and IPTG 0.08 mM to induce sgRNA expression. After a 3-hour incubation with continuous agitation (250 r.p.m.; MaxQ 8000, Thermo Fisher Scientific), suspensions were streaked and incubated overnight at 37°C on LB agar to eliminate pLC10. The next day, single colonies were serially streaked on LB agar supplemented with ERT 0.5 µg/mL, LB agar supplemented with kanamycin (250–512 µg/mL) or apramycin (30-50 µg/mL), and antibiotic-free LB agar. Plates were incubated overnight at 37°C and only colonies able to grow in LB, but not in the other two plates, were selected. Then, pLC10 absence was confirmed by PCR (as described above) and pOXA- 48 absence was determined by PCR using primers targeting two specific different pOXA- 48 conserved regions: (i) the *bla*_OXA-48_ gene (Fw 5’-TTGGTGGCATCGATTATCGG-3’, Rev 5’-GAGCACTTCTTTTGTGATGGC-3’; amplicon size 744 bp) and (ii) *repC* gene (Fw 5’-CGGAACCGACATGTGCCTACT-3’ and Rev 5’-GAACTCCGGCGAAAGACCTTC-3’; amplicon size 852 bp). Confirmed pOXA-48 and pLC10-free colonies were resuspended in LB supplemented with glycerol 13% and stored at -70°C. Additionally, each pOXA-48- carrying/pOXA-48-free bacterial pair was validated with chromogenic medium and antimicrobial susceptibility testing, to determine that the phenotypic profile was consistent with being the same clone with or without pOXA-48 (Table S5). Single colony isolates derived from the curing process were stored. Then, pOXA-48-carrying and pOXA-48-free bacteria were used to inoculate 2 mL LB starter cultures, which were incubated overnight at 37°C with continuous agitation (250 r.p.m.; MaxQ 8000, Thermo Fisher Scientific). Each culture was diluted 1:40 in 2 mL NaCl 0.9% (∼10^7^ colony forming units per millilitre [CFU/mL]) and swabs were used to spread the cell suspension on the Mueller Hinton plates. Antibiotic disks (Biorad) were placed on the plates. Plates were incubated at 37°C for 24 hours and the diameter of the growth inhibition halos were measured. Antibiotic discs tested were: Azythromycin 15 µg (AZM15), Tetracycline 30 µg (TET30), Cefotaxime 30 µg (CTX30), Streptomycin 10 µg (SMN10), Amoxicillin + Clavulanic Acid 20/10 µg (AMC30), Rifampicin 5 µg (RIF5), Fosfomycin 200 µg (FOS200), Chloramphenicol 30 µg (CHL30), Meropenem 10 µg (MEM10).

### Bacterial growth curves

Bacterial cultures were inoculated from freezer stocks in 2 mL of LB and incubated overnight at 37°C with continuous shaking (250 r.p.m.; MaxQ 8000, Thermo Fisher Scientific). Then, each culture was diluted 1:1,000 into fresh LB and was used to fill flat-bottom 96-well plates with 200 µL per well (Thermo Fisher Scientific). Eight replicates from each genotype were included. Then, bacterial cultures were incubated 24 h at 37°C in a plate reader (Synergy HTX Multi-Mode Reader; BioTek Instruments). During incubation, optical densities (OD_600_) were measured after 15 seconds of orbital agitation (282 cpm, 3mm) every 10 minutes. After the incubation, and to discard the potential loss of pOXA-48 during the growth cycles, two replicates of each genotype were serially-diluted in NaCl 0.9% and plated on LB agar plates supplemented with and without amoxicillin + clavulanic acid 250/50 µg/mL (Normon). Agar plates were incubated overnight at 37°C and CFU/mL were estimated. The growth curves data were analysed using Rstudio 2022.12.0+353 (R v4.2.2) and the *flux* v.0.3.0.1 was used to determine the area under the curve (AUC).

### Relative fitness determination by competition assays

To calculate the relative fitness of pOXA-48-carrying isolates compared to their pOXA- 48-free counterparts, competition assays were performed by flow cytometry (CytoFLEX Platform; Beckman Coulter) as in ^16^. Cytometer parameters were: 50 µl min^-1^ flow rate; 22 µm core size; and 10,000 events per well. *E.coli* J53^35^ (a sodium azide resistant mutant of *E. coli* K-12) carrying the pBGC plasmid (accession number MT702881, Figure S1C) was used as a common competitor in all competition assays. pBGC is a non-mobilizable plasmid that contains the *gfp* gene under the control of the P*_BAD_* promoter. Hence, GFP production is induced by the presence of arabinose. Two sets of competitions were performed for each isolate: pOXA-48-free *vs*. *E. coli* J53/pBGC, and pOXA-48-carrying *vs*. *E. coli* J53/pBGC with six replicates per competition. Initial pre-cultures were inoculated from freezer stocks in 200 µL of LB and incubated overnight with continuous shaking 225 r.p.m. at 37°C in 96-well plates (Thermo Fisher Scientific). Next day, bacterial cultures were mixed 1:1 and diluted 10,000-fold in fresh LB. Then, mixtures were competed for 22 h in LB at 37°C and 225 r.p.m. To determine the initial proportions, initial mixes were diluted 2,000-fold in 200 µL of NaCl 0.9% with L-arabinose (Sigma-Aldrich) 0.1%, and incubated at 37°C at 225 r.p.m. during 1.5 hours to induce GFP expression before performing flow cytometry determinations. After 22 h of incubation, final proportions were again determined after a 2,000-fold dilution of the cultures in the same conditions as before. The fitness of each strain relative to the same common competitor —*E. coli* J53/pBGC— was determined using the formula (1):

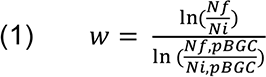

where *w* is the relative fitness of the pOXA-48-carrying or pOXA-48-free isolates compared to the *E. coli* J53/pBGC, *Ni* and *Nf* are the number of cells of the pBGC-free clone at the beginning and end of the competition, and *Ni*,pBGC and *Nf*,pBGC are the number of cells of the pBGC-carrying *E. coli* J53 at the beginning and end of the competition, respectively. The fitness of the pOXA-48-carrying parental isolates relative to the pOXA-48-free cured isolates (*w_p_*) was calculated using the average result of six independent replicates of each competition and using the formula (2):

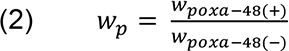

where *w*pOXA-48(+) corresponds to the relative fitness of the pOXA-48-carrying clone and *w*pOXA-48(-) to the relative fitness pOXA-48-free clone, both compared to the common *E. coli* J53/pBGC competitor. The error propagation method was used to calculate the standard error of the resulting value. Additionally, *gfp* induction was monitored in each competition by growing individual *E. coli* J53/pBGC cultures. Potential plasmid loss during the competition was discarded by controlling for plasmid loss during growth curves (Figure S3). Importantly, controls to discard for potential conjugative transfer of pOXA-48 during competitions were performed by plating the final time points of two replicates of each competition assay on ertapenem (0.5 µg/mL), chloramphenicol (200 µg/mL) and sodium azide (100 µg/mL). No transconjugants were detected, probably due to the small initial population size of the competing populations and the strong shaking regime.

### Genomic DNA extraction and sequencing

Genomic DNA (gDNA) was extracted using the Wizard gDNA purification kit (Promega Corporation). From the 35 wild-type strains that were cured from pOXA-48, a subset of 22 strains was sequenced at SeqCoast Genomics (https://seqcoast.com/), using the NextSeq 2000 platform (coverage > 60×). Sequencing data is available under the BioProject PRJNA962867. For the remaining 13 wild-type strains, we used the Illumina data generated previously^21^ (BioProject PRJNA626430). The genomes of six wild-type strains were also sequenced with long reads to generate closed references, either with Nanopore technology at the Microbial Genome Sequencing Center (MiGS) or with PacBio RSII at The Norwegian Sequencing Centre (see Table S1). Long-read data and closed assemblies were submitted to SRA and GenBank under the BioProject PRJNA626430. To control for putative curing-derived off-target mutations, a subset of nine cured strains was sequenced at the Wellcome Trust Centre for Human Genetics (Oxford, UK), using the Illumina HiSeq4000 platform, or the Microbial Genome Sequencing Center (MiGS), using the NextSeq 2000 platform (Table S1; data available at BioProject PRJNA962867). The closed sequences of three wild-type strains were retrieved from PRJNA838107^20^ (Table S1).

### Bioinformatic analyses

A detailed description of bioinformatics methods is provided at https://github.com/LaboraTORIbio/CRISPR_cured_pOXA-48. Illumina reads were trimmed with Trim Galore v0.6.4 (options --quality 20 --length 50 --illumina --paired, https://github.com/FelixKrueger/TrimGalore) or Trimmomatic^36^ v0.39, using the maximum information quality filtering algorithm (MAXINFO:50:0.8) and removing adapters. Hybrid assemblies of the wild-type strains C325, CF12, CF13, H53, J57 and K147 were obtained with Unicycler^37^ v0.4.9. *De novo* genome assemblies were generated with SPAdes^38^ v3.15.2 (options --isolate --cov-cutoff auto). Assembly quality was assessed with Bandage^39^ v0.8.1 or QUAST^40^ v5.0.2. Complete genomes were annotated with PGAP^41^ v2021-07-01.build5508 and draft assemblies with Prokka^42^ v1.14.6. Multilocus Sequence Type was assigned with MLST v2.21.0 (https://github.com/tseemann/mlst).

Phylogenetic trees based on mash distance were constructed with mashtree^43^ v1.2.0 using the whole-genome assemblies and a bootstrap of 100.

Variant calling was performed with breseq^44^ v0.35.6 or v0.35.7. First, to discard false positive calls due to misassemblies, the Illumina reads of the wild-type strains were mapped to their respective closed genomes. Then, the Illumina reads of the cured strains were mapped to their corresponding wild-type reference to identify mutations in cured strains. To further confirm or discard confusing mutations, the Illumina reads of the wild-type strains were mapped to the draft genomes of the cured strains. All mutations reported by breseq —single nucleotide polymorphisms (SNPs), missing coverage (deletions) and new junction (genomic rearrangements) evidences— were analysed.

The identified mutations were further characterised with different methods depending on their nature. Functional effects of missense variants were predicted with the SNAP2^45^ webserver (https://rostlab.org/services/snap2web/), where scores lower than 0 indicate neutral effects on protein function. Synonymous variants were not investigated since all would be predicted as neutral. PSIPRED^46^ v4.0 was used to predict changes in protein secondary structure caused by frameshift variants, and domain information was obtained from InterProScan^47^ 5. The nucleotide sequences of the affected intergenic regions were scanned for promoters using phiSITE’s PromoterHunter^48, 49^. The nucleotide sequences of the mutated genes and intergenic regions were submitted to a BLASTn^49^ search against the NCBI database (nr/nt collection, restricted to Enterobacteria (taxid:543)) to determine if the mutations are present in other enterobacteria.

To determine whether the identified variants could be due to off-target cuts of the CRISPR-Cas9 system, the nucleotide sequences of the guides sgOXA48 and sgPEMK were aligned to a 40 bp subregion enclosing the mutations using EMBOSS Needle^50^ v6.6.0.0. This tool finds the best global alignment between two sequences. Therefore, if mutations were produced by the CRISPR-Cas9 system, Needle would show the sgRNA alignment that most probably caused the off-target cleavage.

The selective elimination of plasmid pOXA-48 from all cured strains was confirmed by inspecting the missing coverage evidence reported by breseq and by identifying plasmid replicons with ABRicate v1.0.1 (https://github.com/tseemann/abricate), using the PlasmidFinder database.

### Statistical analysis

Statistical analyses were performed in RStudio (R v4.2.2) with R base and packages *tidyverse*, *outliers*, *ggplot2* and *ggpubr*. To test homoscedasticity and normality, Bartlett and Shapiro-Wilk tests were performed. Then, according to each data structure, parametric and non-parametric tests were performed (see main manuscript for each test). To compare the distributions of fitness effects between our collection and Alonso-del Valle *et al*.’s 2021 collection, different tests were performed. Data in our collection did not follow a normal distribution (Shapiro-Wilk test, *P* = 7.36 x 10^-6^), since strain AJ_N46 is an outlier (Grubbs test, *P* = 1.07 x 10^-6^), although both data distributions did not differ according to a Kolmogorov-Smirnov test, (*D* = 0.17, *P* = 0.56). When removing the outlier, data distribution was normal (Shapiro-Wilk test, *P* = 0.095). In addition, the means of both distributions were not different as reported by a two-sample *t* test (*t* = 0.23, df = 77.67, *P* = 0.83) and Wilcoxon rank sum test (*W* = 923, *P* = 0.6715).

## Data availability

The sequencing data supporting the findings of this study are available at the National Center for Biotechnology Information Database with accession number PRJNA962867.

The closed genomes of the six wild-type strains can be accessed from PRJNA626430.

The raw experimental data obtained in this study are available as Source Data. The remaining R-GNOSIS sequences can be found in ^21^.

## Code availability

The code generated during the study can be found at https://github.com/LaboraTORIbio/CRISPR_cured_pOXA-48.

## Author contributions

A.S.M. and J.D.F. conceptualised the study. A.F.-C., A.A.V., P.V.-C. and J.D.F. designed the methodology and performed the experiments. L.T.-C, J.S.-D., A.F.-C, J.D.F. and A.S.M. contributed to data analyses. A.S.M. was responsible for funding acquisition, supervision and prepared the original draft of the manuscript. J.D.F. undertook the reviewing and editing process. All authors supervised and approved the final version of the manuscript.

## Acknowledgements

We thank the R-GNOSIS group, the volunteers and the medical staff from the Hospital Universitario Ramón y Cajal (Madrid, Spain) involved in the sample isolation process.

## Supplementary Material

### Supplemental Results

#### Analysis of mutations affecting coding regions in cured clones

The genomes of the cured strains presented in total eight-point mutations in coding genes (Table S3). Four SNPs were predicted to have neutral effects on protein function: a synonymous variant in the iron donor protein CyaY in C288c2, a synonymous variant affecting the putrescine ABC transporter ATP-binding subunit PotG in C325c1, and two missense variants affecting a phage capsid protein in CF12c1 (SNAP2 score -88, accuracy 93%) and the type VI secretion system baseplate subunit TssF in H53c1 (SNAP2 score -64, accuracy 82%). It is worth noting that these mutations could indirectly impact protein function by altering gene expression^1, 2^. The remaining four SNPs were predicted to alter protein function. The frameshift mutation affecting gene *pdeR* in CF12c1 compromises protein secondary structure (Figure S6), producing the elongation of the C-terminus (+22 amino acids), which comprises the EAL domain of PdeR. Mutations in this protein, a member of the curli fimbrae biosynthesis cascade, are related to increased biofilm formation^3^. Another mutation restored the reading frame of the frameshifted *znuB* gene in H53c1, a subunit of the zinc ABC transporter complex. The third variant introduced a premature stop codon that results in the loss of more than a third of an integrase domain-containing protein in strain K147c1. The loss of function of this protein would probably only affect integron biology, impeding its excision. Thus, it should not produce large-scale physiological changes in the cured strain with respect to the wild-type strain. The fourth SNP also introduced a premature stop codon in the *ompC* gene of C288c2. This resulted in a truncated version of the protein, cutting down from 367 to 171 amino acids. The nucleotide sequences of the mutated genes were submitted to a BLASTn search against the NCBI database. In C288c2, the synonymous SNP in the *cyaY* gene was also found in other enterobacteria (e.g. CP077379.1). In the case of H53c1, SNPs present in both *znuB* (as seen earlier, Figure S7) and *tssF* were also present in other enterobacteria (e.g. CP043597.1 and CP081896.1, respectively), suggesting these are naturally-occurring variants. Lastly, we detected two probable insertion sequence (IS) rearrangements. One of them was a possible event of an IS jump from the IncF plasmid of J57c1 to its chromosome, inserting into the yhjQ gene and probably producing a knock-out. This gene is involved in the biosynthesis of cellulose, and its disruption can reduce bacterial aggregation and biofilm formation4. The other one was observed in K153c2: an IS1 element from its chromosome was integrated within a coding region of a putative AAA family ATPase, which are involved in a wide variety of functions, potentially producing a knock-out.

#### Analysis of mutations affecting intergenic regions in cured clones

We analysed if the observed intergenic SNPs could be located within putative promoters. In the case of CF12c1, the SNP located 50 bp upstream *glpA* did not apparently fall within predicted promoters in the sense strand that could control the expression of *glpA* and downstream genes (Figure S8A). Still, this mutation could affect other regulatory regions or the transcription of non-coding RNAs. In J57c1, the SNP located 94 bp upstream the catecholate siderophore receptor *fiu* gene is positioned within a predicted promoter in the sense strand (final score 2.72) and near another promoter with higher score (2.79) (Figure S8B). Therefore, the expression of the *fiu* gene and downstream genes could be altered. In the IncF plasmid of J57c1 there are multiple intergenic mutations between positions 90533-90554 bp that could be due to the possible excision and insertion of the IS element into the chromosome of J57c1. These SNPs fall near predicted promoters for the sense strand, and thus, could affect transcription of the ISNCY-like element ISKpn21 family transposase (Figure S8C). Nonetheless, these mutations had low read coverage and alignment quality (multiple contiguous SNPs), and thus could represent false positive calls. In K153c2, another IS1 element was observed integrated into an intergenic region between the *fyuA* gene and a hypothetical protein, 77 bp upstream the latter. As with mutations in coding regions, the sequences of the mutated intergenic regions were also submitted to a BLASTn search against the NCBI database. However, we could not find the observed mutations in other organisms.

#### Analysis of possible off-target events

We aligned the sequences of the single-guide RNAs (sgOXA48 and sgPemK) to the regions surrounding the identified SNPs (see Methods). To recognize and produce off-target cuts, the alignment of the sgRNA must have less than eight mismatches, short or no gaps, and more importantly, the Cas9 protospacer adjacent motif (PAM) sequence NGG or NAG at the 3’ of the sgRNA^5^. Most alignments had gaps and/or more than seven mismatches and lacked the PAM sequence. Only two alignments had less than eight mismatches and presented the PAM sequence NGG, constituting possible off-target events (Figure S9). First, sgOXA48 aligned against the region encompassing the *znuB* mutation in H53 —that restores the reading frame of *znuB* in H53c1— with only five mismatches (Figure S9A). However, the mutation affecting *znuB* in H53c1 is a naturally-occurring variant. In fact, we could not find any enterobacteria in the NCBI database that carried the frameshift variant of H53. Second, the sgPemK guide aligned against the region surrounding the *ompC* mutation in C288 (Figure S9B). However, the combination of gaps and/or mismatches located within the PAM-proximal region of the sgRNAs would most probably impede Cas9 cleavage^6^. Moreover, none of the other analysed strains showed mutations in the *ompC* nor the *znuB* genes, contradicting the possibility of off-target CRISPR-Cas9 activity.

## Supplementary Figures

**Figure S1.**
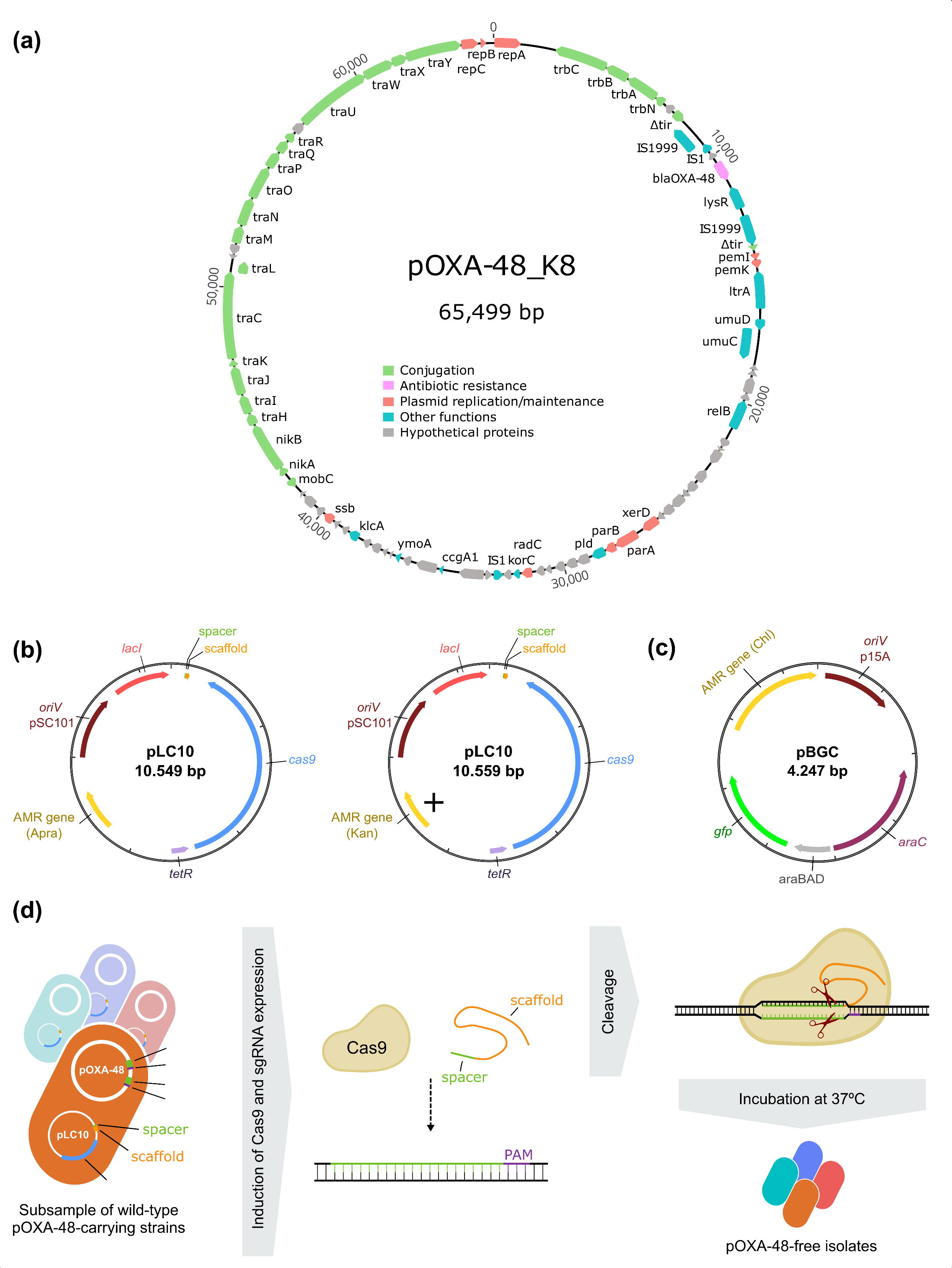
Plasmids used in this study and plasmid pOXA-48 curing protocol. (a) Genetic map of pOXA-48_K8 (accession number MT441554), the most prevalent plasmid variant in our collection. Open reading frames are represented with arrows. Colours indicate gene functions (see legend). (b) Genetic maps of pLC10 plasmids, encoding the CRISPR-Cas9 system and a gene conferring resistance to apramycin (left) or kanamycin (right). The *cas9* gene and the single-guide RNA (sgRNA: spacer + scaffold) are under the control of the anhydrotetracycline (ATC)-inducible and isopropyl β-d-1-thiogalactopyranoside (IPTG)-inducible promoters P*_tet_*and P*_lac_*, respectively. pLC10 also encodes a thermosensitive replication initiation protein (pSC101-based) that is not functional at 37°C. (c) Genetic map of pBGC (accession number MT702881), used in competition assays. (d) Experimental design of pOXA-48 curing protocol (see Methods). Plasmid pLC10, encoding the sgRNA targeting either the *bla*_OXA-48_ or *pemK* pOXA-48 genes (see PAM sequence 5’-NGG-3’ in pOXA-48), was transformed into a subsample of wild-type pOXA-48-carrying enterobacterial strains. The expression of the *cas9* gene and the sgRNA was induced with ATC and IPTG, respectively, allowing cleavage of pOXA-48 at either target. Plasmid pLC10 was eliminated by incubating at 37°C. pOXA-48-free strains were selected from colonies growing only in LB agar plates without antibiotics.

**Figure S2.**
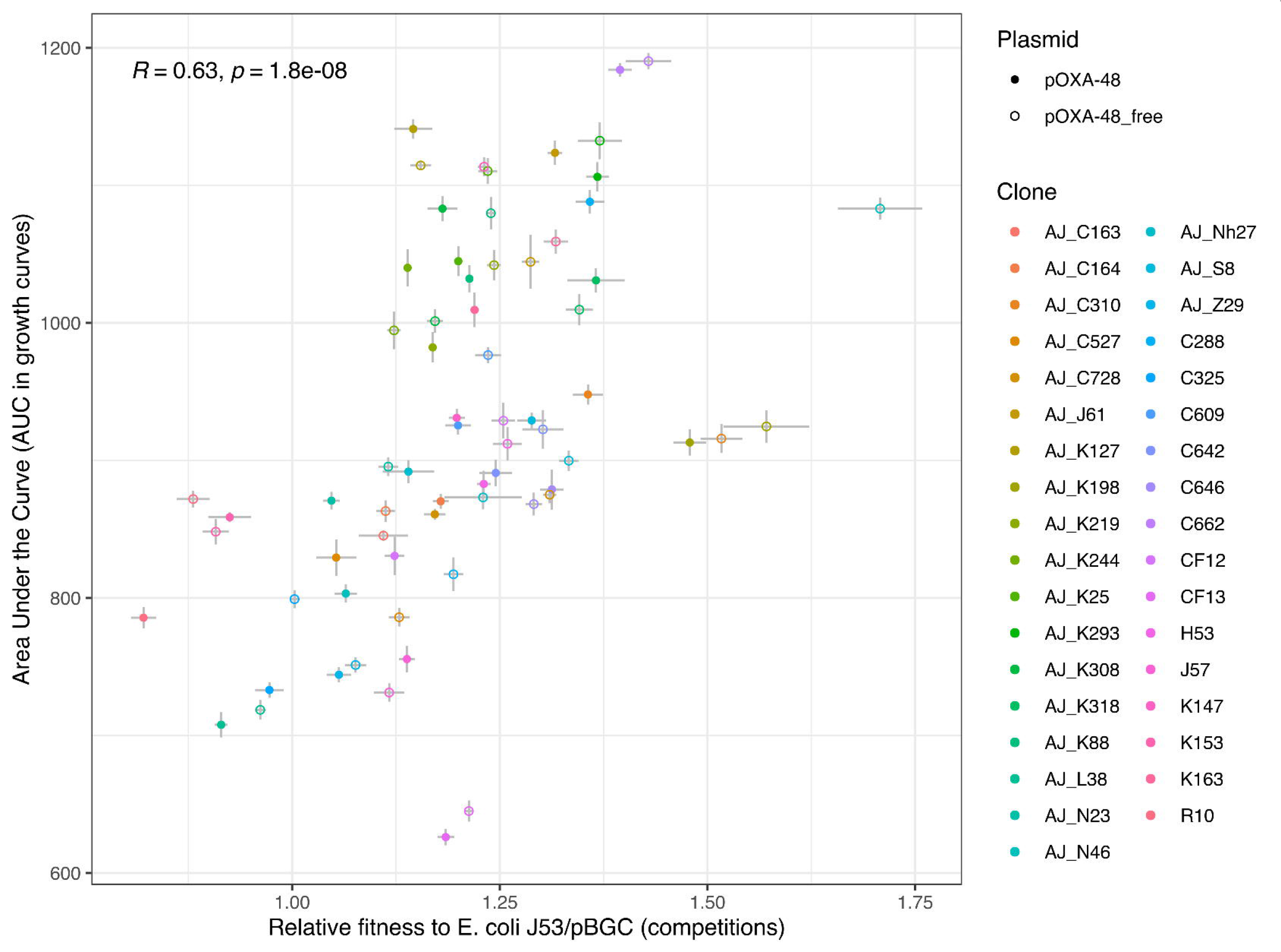
Correlation between the relative fitness and the area under the growth curves of each pOXA-48-carrying and pOXA-48-free clones. Note that the relative fitness of each clone was obtained by competing each pOXA-48-carrying and pOXA-48-free clone against a common competitor (*Escherichia coli* J53/pBGC). Each dot indicates the mean value (n = 6 for competition assays and n = 8 for growth curves). Lines indicate the standard error of the mean. Clones are indicated by colours and the presence/absence of pOXA-48 is indicated by full or empty circles. The Spearman’s rank correlation between the fitness of each clone relative to *E. coli* J53/pBGC and the area under the growth curve is indicated in the top-left section of the figure (Spearman’s rank correlation *rho =* 0.626, *S* = 20446, *P* = 1.84 x 10^-8^).

**Figure S3.**
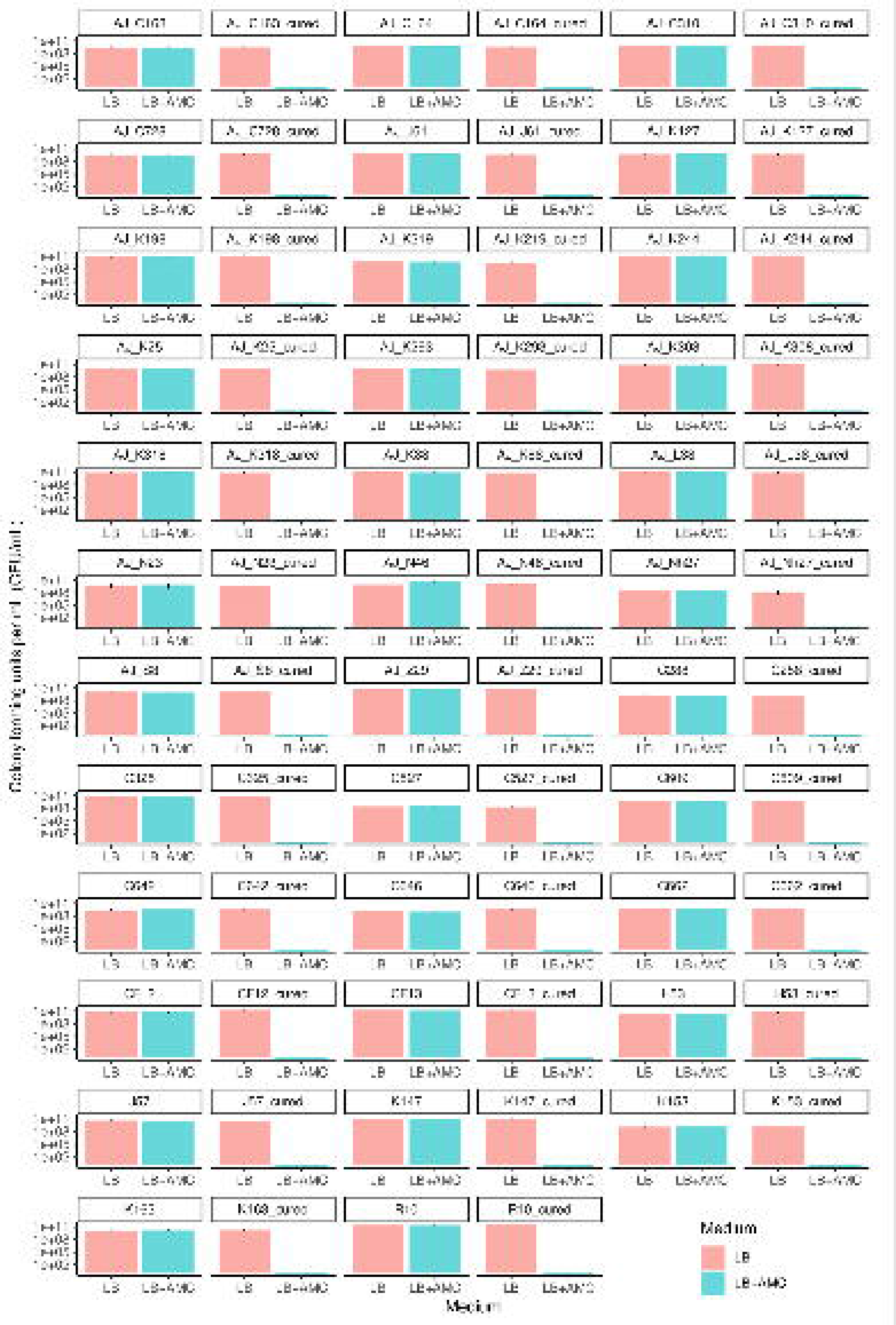
Measuring pOXA-48 loss during growth cycles. After growth curves, cells were serially-diluted and plated on agar plates supplemented with and without pOXA-48 selective antibiotics (AMC, Amoxicillin + clavulanic acid). Then, colony forming units (CFU/mL) were estimated (see Methods). Colours indicate each medium, bars indicate the CFU/mL and the error bars indicate the standard deviation of the mean. Note that (i) in cured strains no colony forming units were detected in AMC treatments and (ii) there are no differences between CFU on selective and non-selective media for pOXA-48- carrying clones (Kruskal-Wallis, *chi-squared* = 0.00028, *P* = 0.9867), indicating that plasmid pOXA-48 was maintained during the growth cycle.

**Figure S4.**
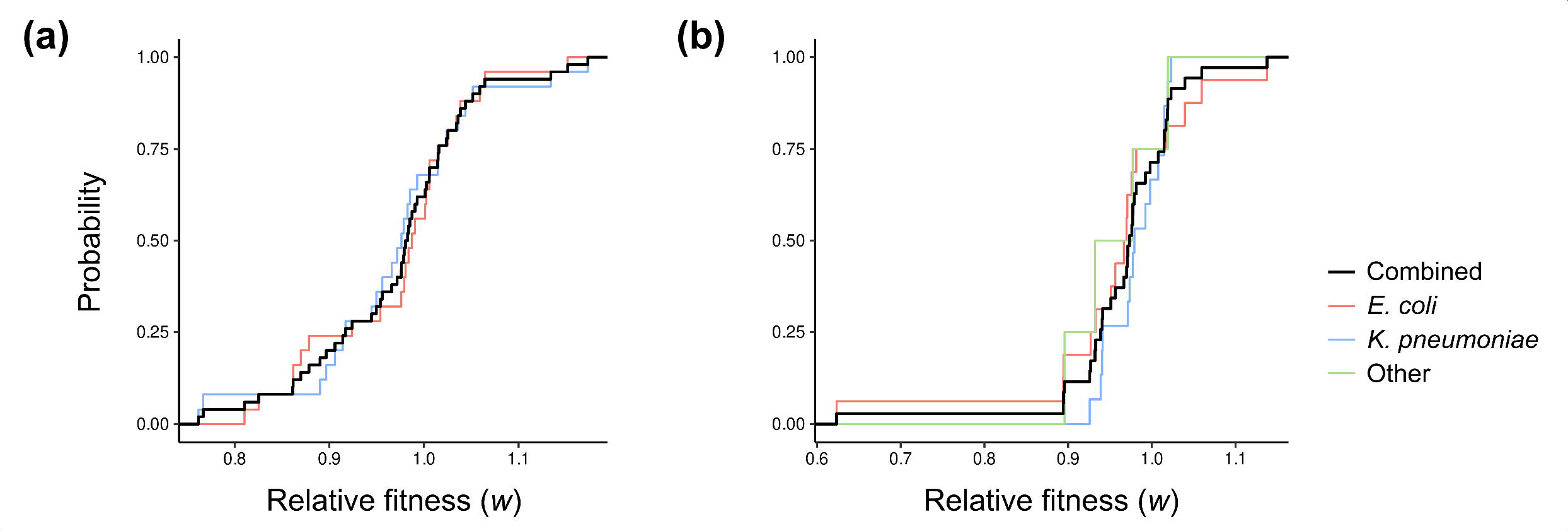
Cumulative distribution functions (CDF) of relative fitness (*w*) of pOXA-48- carrying strains. (a) CDF of the collection from Alonso-del Valle *et al*. 2021, which includes naive, ecologically compatible, pOXA-48-carrying enterobacteria (n = 50). (b) CDF of the collection of wild-type pOXA-48-carrying enterobacteria analysed in this work (n = 35). Lines indicate the CDF of all combined strains (black line) and strains separated by species (red line for *E. coli*, blue line for *K. pneumoniae* and green line for other species).

**Figure S5.**
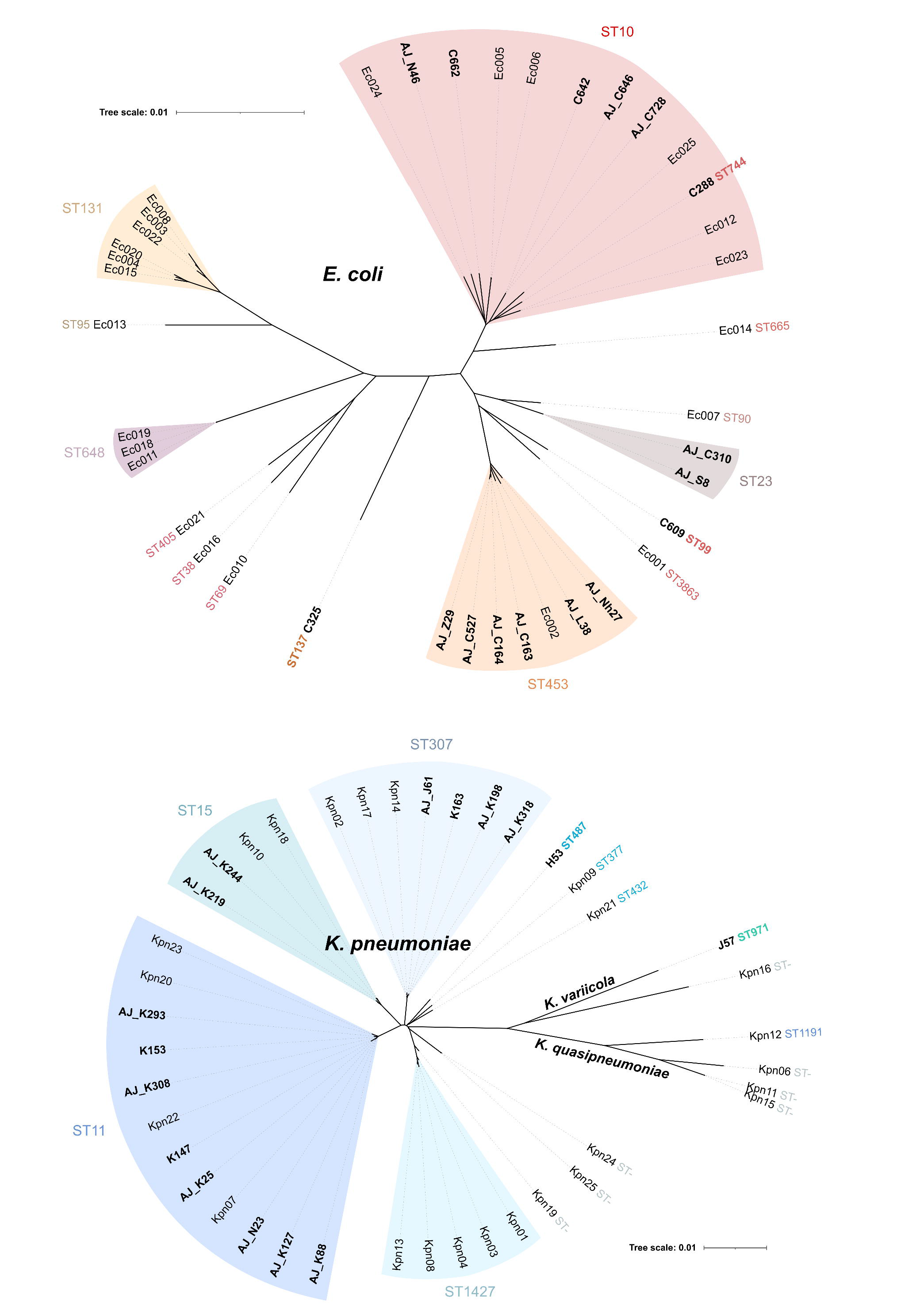
Unrooted phylogenetic trees of *Escherichia coli* (upper panel) and *Klebsiella* spp. (lower panel) strains from the collections of pOXA-48-cured wild-type enterobacteria used in this study and naive, ecologically compatible pOXA-48 carriers^7^. Phylogenies were constructed from mash distances between whole-genome assemblies, represented by branch lengths. Multilocus Sequence Type (ST) is indicated.

**Figure S6.**
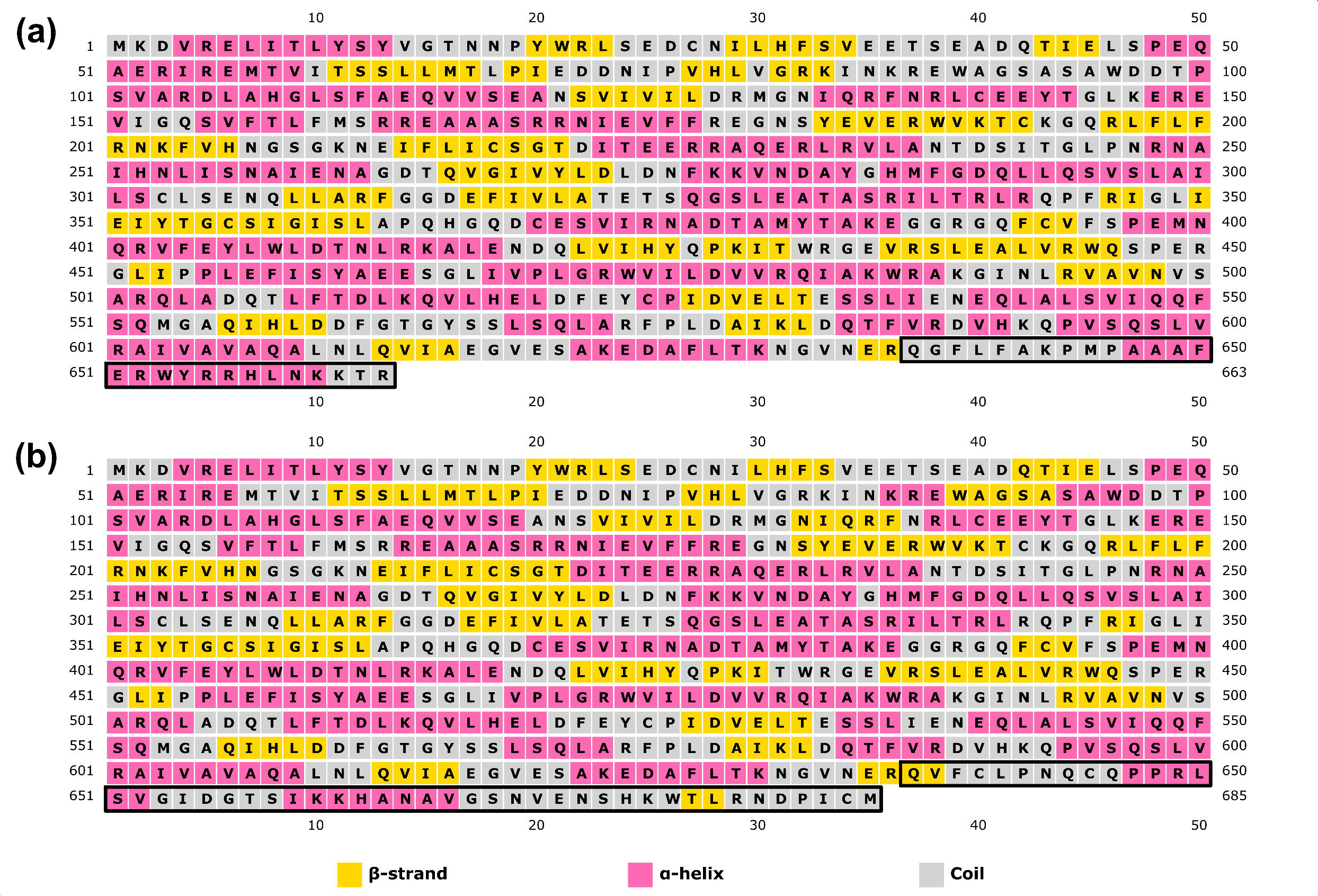
Secondary structure prediction by PSIPRED of PdeR. The PdeR protein of the wild-type CF12 strain has 663 amino acids (a). The frameshift variant of the cured strain affecting the residue Q638 of PdeR produces an elongation of +22 amino acids in the C-terminus that causes a change in secondary structure (b). The reference (a) and affected regions (b) are indicated by a black box.

**Figure S7.**
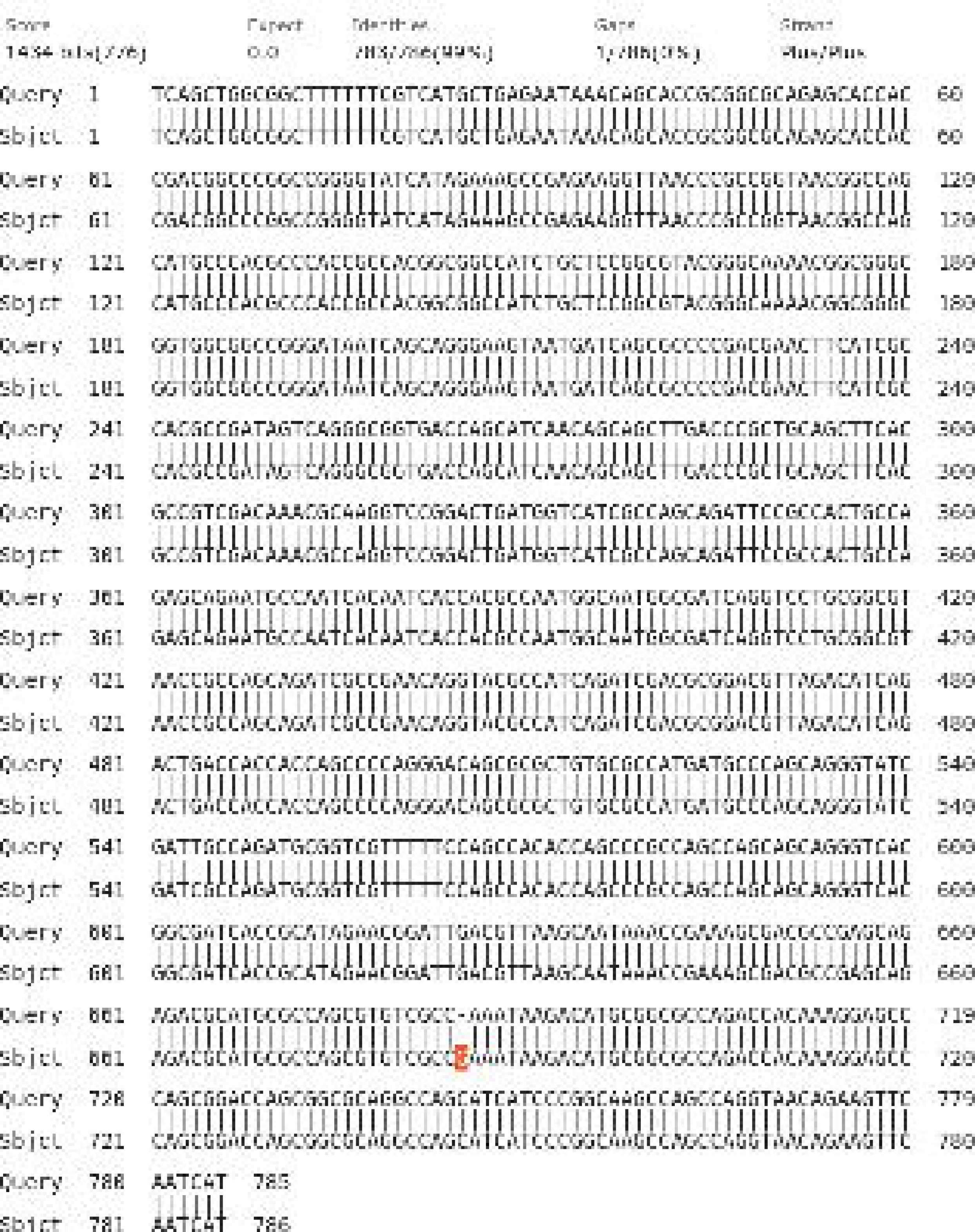
BLASTn alignment of H53’s *znuB* (Query) and K147’s *znuB* (Sbjct). The +G in K147 (also present in H53c1 and in other enterobacteria of the NCBI database), that restores the reading frame of *znuB*, is highlighted in orange.

**Figure S8.**
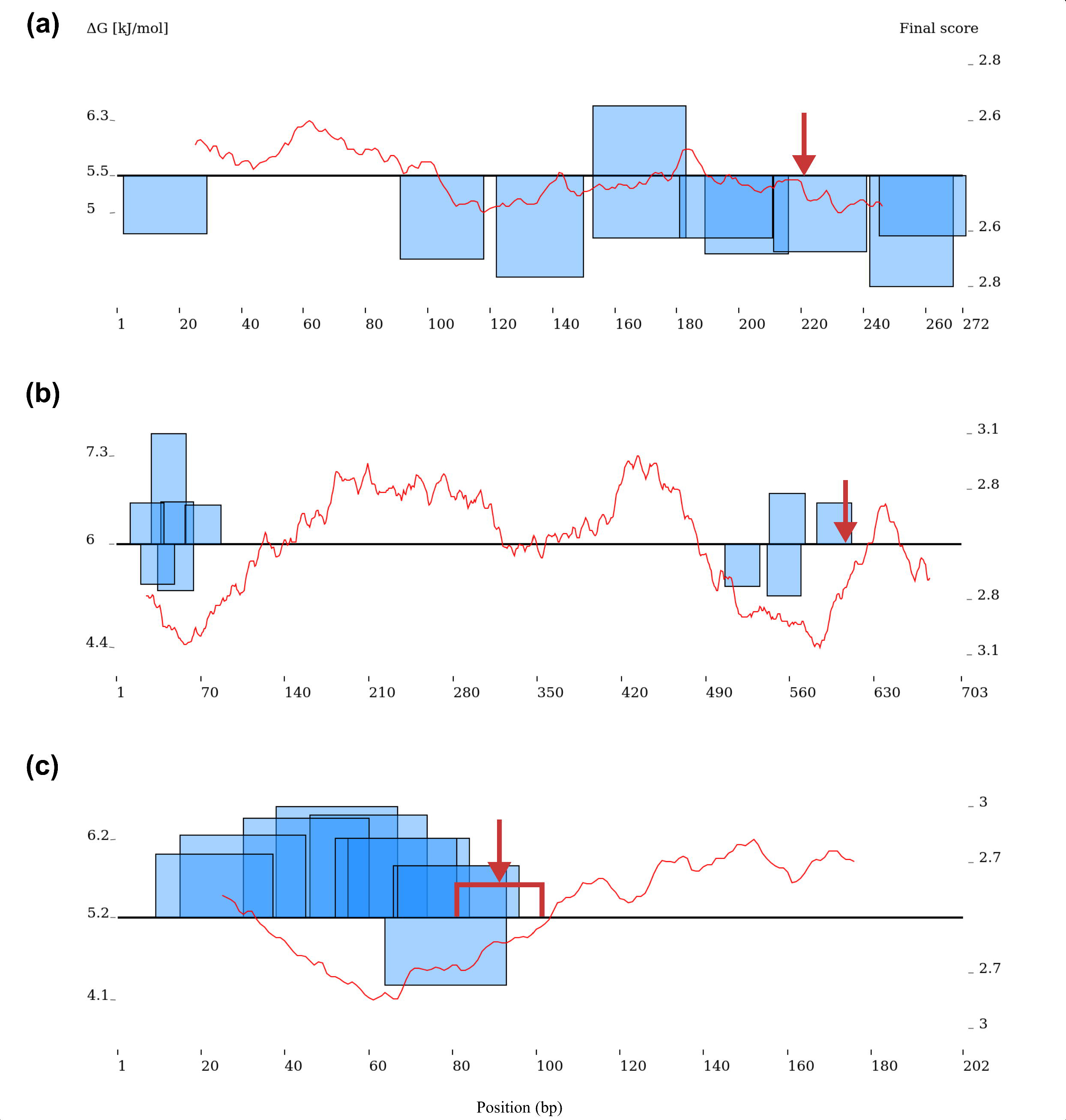
Putative promoters predicted by PromoterHunter in the mutated intergenic regions. (a) Intergenic region between genes *glpT* (antisense strand) and *glpA* (sense strand) in CF12. (b) Intergenic region between genes encoding the zinc-binding alcohol dehydrogenase family protein (sense) and the catecholate siderophore receptor Fiu (sense) in J57. (c) Intergenic regions between the genes encoding the GNAT family N-acetyltransferase (sense) and the ISNCY-like element ISKpn21 family transposase (sense) in the IncF plasmid of J57. In (a) and (b) the location of the SNPs is indicated with a red arrow. In (c), a red bracket also indicates the borders of the highly mutated region. Blue boxes above the horizontal black line represent putative promoters in the sense strand; boxes below the horizontal line represent putative promoters in the antisense strand. Left *y* axis and the red line represent the Gibbs free energy (ΔG) distribution along the sequence. Right *y* axis and the height of the blue boxes indicate the final scores of the putative promoters.

**Figure S9.**
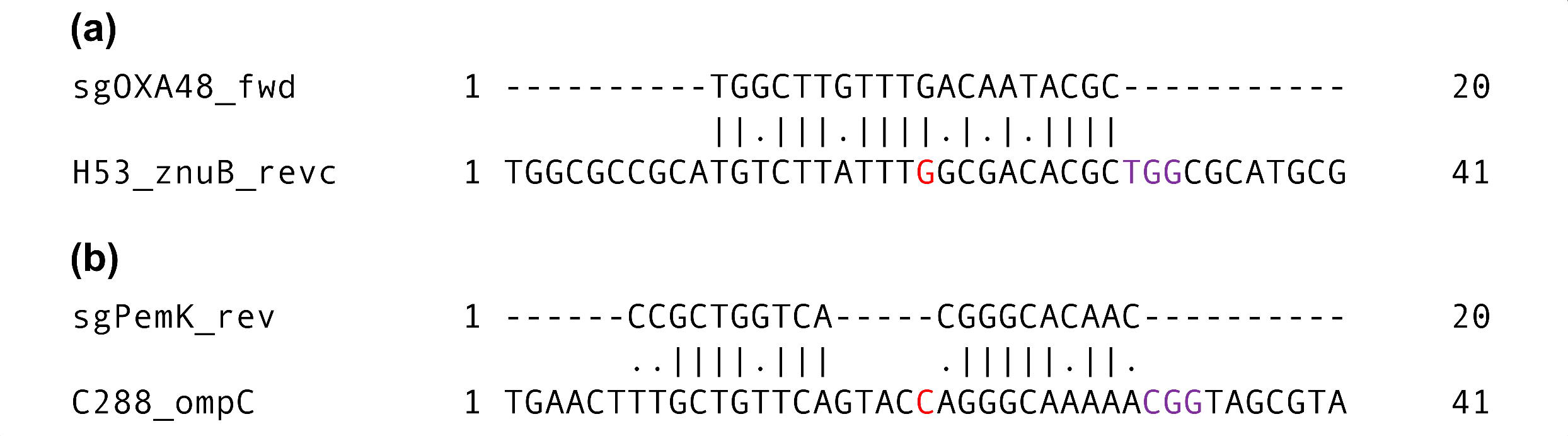
Global alignments of sgRNAs against the regions enclosing mutations in H53’s *znuB* (a) and C288’s *ompC* (b). Only these alignments presented less than eight mismatches and the PAM sequence (NGG). The SNP position is indicated in red and the PAM sequence in purple. Mismatches are represented with dots. Fwd: forward sgRNA, rev: reverse sgRNA; revc: reverse complementary DNA sequence.

